# Mid-infrared photoacoustic brain imaging enabled by cascaded gas-filled hollow-core fiber lasers

**DOI:** 10.1101/2024.04.02.587715

**Authors:** Cuiling Zhang, Kunyang Sui, Marcello Meneghetti, Jose Enrique Antonio-Lopez, Manoj K. Dasa, Rune W. Berg, Rodrigo Amezcua-Correa, Yazhou Wang, Christos Markos

## Abstract

**Significance:** Extending the photoacoustic microscopy (PAM) into the mid-infrared (MIR) molecular fingerprint region constitutes a promising route towards label-free imaging of biological molecular structures. Realizing this objective requires a high-energy nano-second MIR laser source. However, existing MIR laser technologies are limited to either low pulse energy or free-space structure which is sensitive to environmental conditions. Fiber lasers are promising technologies for PAM for their potential of offering both high pulse energy and robust performance against environmental conditions. However, MIR high energy fiber laser has not yet been used for PAM because it is still at the infant research stage.

**Aim:** We aim to employ the emerging gas-filled anti-resonant hollow-core fiber (ARHCF) laser technology for MIR-PAM for the purpose of imaging myelin-rich regions in a mouse brain.

**Approach:** This laser source is developed with a ∼2.75 μJ high-pulse-energy nano-second laser at 3.4 μm, targeting the main absorption band of myelin sheaths, the primary chemical component of axons in the central nervous system. The laser mechanism relies on two-orders gas-induced vibrational stimulated Raman scattering (SRS) for nonlinear wavelength conversion, starting from a 1060 nm pump laser to 1409 nm through the 1^st^ order Stokes generation in the nitrogen-filled 1^st^ stage ARHCF, then, from 1409 nm to 3.4 μm through the 2^nd^ stage hydrogen-filled ARHCF.

**Results:** The developed Raman laser was used for the first time for transmission-mode MIR-PAM of mouse brain regions containing rich myelin structures.

**Conclusions:** This work pioneers the potential use of high-energy and nano-second gas-filled ARHCF laser source to MIR-PAM, with a first attempt to report this kind of fiber laser source for PAM of lipid-rich myelin regions in a mouse brain. The proposed ARHCF laser technology is also expected to generate high-energy pulses at the ultraviolet (UV) region, which can significantly improve the lateral resolution of the PAM.

## 1 Introduction

Photoacoustic microscopy (PAM) is a prominent non-invasive imaging modality that uses optical excitation to generate ultrasound signals, allowing the visualization of biomedical tissues, inorganic materials, or complex samples with larger penetration depths when compared to other optical imaging techniques [1–4]. In recent years, PAM within the visible and near-infrared (NIR) wavelength regions have unveiled a wealth of functional information, contributing to advancements in various research fields [5–9]. Currently, the scientific community focuses on moving PAM technology to the mid-infrared (MIR) wavelength domain, opening the door to new opportunities in microscopy with emphasis on probing specific molecular vibrational bands, such as those associated with CH_2_ groups [10–15]. For instance, He *et*.*al*. reported the use of mid-infrared photoacoustic microscopy (MIR-PAM) for mapping the lipid compositions (CH_2_ stretching transition) in mouse brain and kidney tissue at ∼3.4 μm [10]. Furthermore, MIR-PAM enabled the imaging of carbohydrates (at ∼9.2 μm), lipids (at ∼3.5 μm), and proteins (at ∼6.4 μm) in living cells and tissues [11,12]. Compared with the NIR-PAM, the lateral resolution of the MIR-PAM is diffraction-limited to the long wavelength of the MIR laser. This challenge was recently addressed by adding an ultraviolet (UV) pulsed laser as a probe, to enhance the PAM resolution down to the nanometer scale to image lipids, proteins, and nucleic acids [13].

Quantum-cascaded lasers (QCLs) and optical parametric oscillators (OPOs) are the main light sources that have been used for MIR-PAM because mainly of their broad wavelength selectivity [10–15]. However, QCL technology is limited by low peak power of only few watts, leading to a low pulse energy of only few nanojoule at even tens of nanosecond pulse duration [11,14-15]. Some efforts have been made to circumvent this issue, including mitigating the laser energy attenuation caused by the ambient gas absorption such as creating a nitrogen (N_2_)-filled atmosphere, and prolonging the pulse duration to tens of nanosecond. Based on these methods, QCL has been used for scanning-mode-based PAM, however, the pulse energy is too low to be used for real-time PA modalities [14-16]. OPO technology, on the other hand, can deliver high pulse energy of tens microjoule. However, its free-space structure [10] introduces practical limitations due to its sensitivity on environmental conditions such as vibrations, humidity, and temperature.

Fiber lasers are promising alternatives for PA technology because they have the potential to deliver both high pulse energy and robust performance against environmental conditions. Driven by these advantages, mature visible and NIR fiber laser technologies have been recently proposed for PA technology [17-20]. However, MIR high-energy fiber lasers have not yet been used for PAM. In this work, we developed a novel MIR fiber laser technology for PA imaging for the first time. Specifically, the fiber laser emits high energy nanosecond pulses at the absorption peak of myelin at 3.4 μm wavelength to image the neurons in the brain slice [10,12]. The mechanism of the proposed laser is based on the stimulated Raman scattering (SRS) in gas-filled anti-resonant hollow-core fiber (ARHCF), which is an emerging fiber technology, that allows confinement of the laser beam within its hollow region resulting in very strong light-gas (atomic or molecular) interactions (thus efficient SRS frequency) [21,22]. This property not only enables the generation of strong pulses over a broad wavelength range from ultraviolet (UV) to MIR (as summarized in Table S1) [23–28], but also is not limited from the damage threshold of the glass. Therefore, compared to other nonlinear media, gases are considered, in a sense, self-healing, unlike solids which can become permanently damaged [22] and thus allowing energy and intensity scaling of nonlinear devices to their most extreme limit [29]. This highlights its potential to broaden the spectral range beyond the existing fiber lasers utilized in PAM, such as standard doped silica fibers [17-19].

## 2 Material and methods

Figure 1 shows the configuration of the entire system combining the proposed MIR gas-filled fiber laser source and the PAM.

**Fig. 1.**
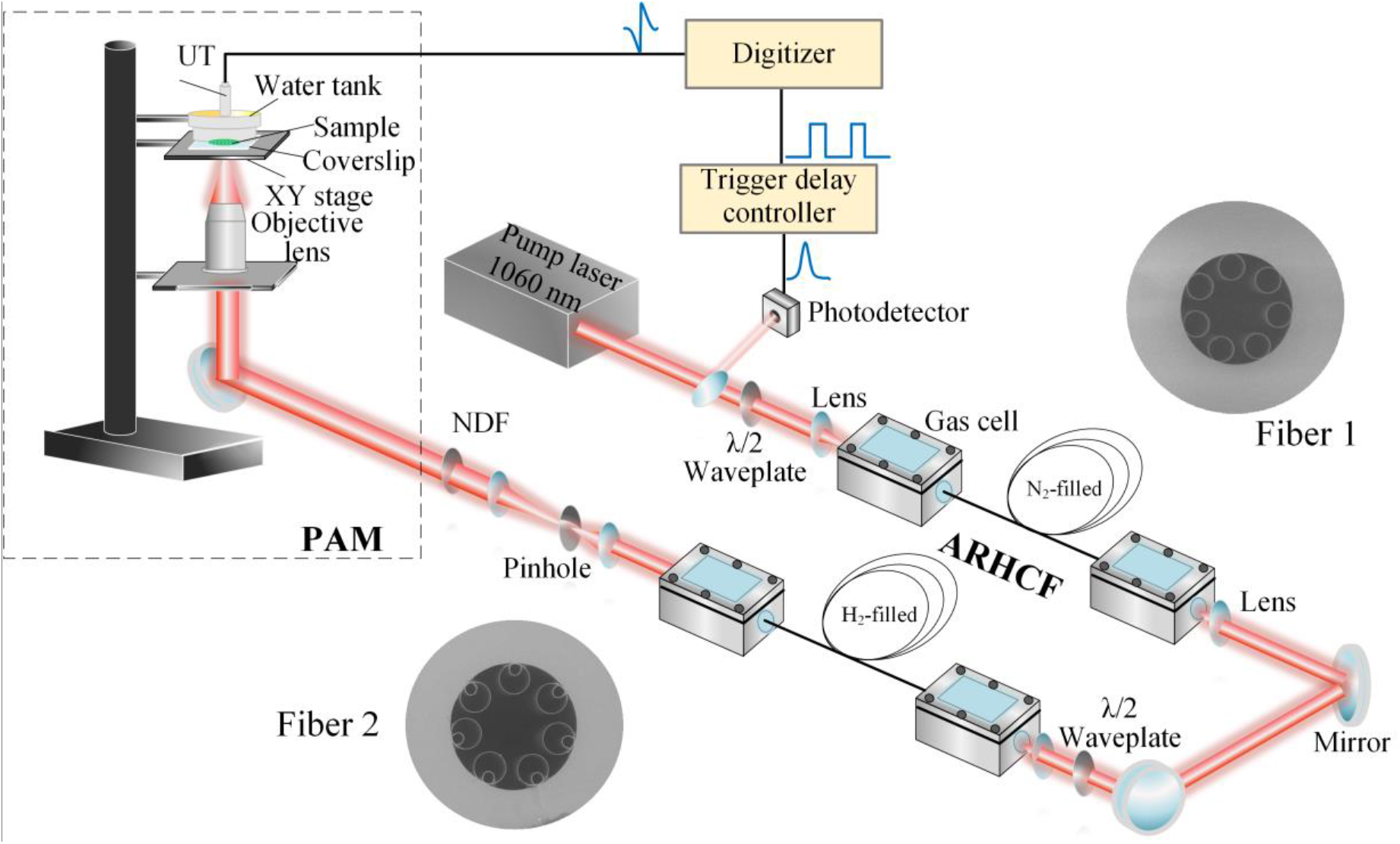
Overall MIR-PAM system. The system consists of two-stage cascaded gas-filled ARHCF laser followed by a transmission mode PAM. Fiber 1 and Fiber 2: SEM images of the 1st and 2nd stage ARHCFs, respectively. ARHCF: anti-resonant hollow-core fiber. NDF: neutral density filter. UT: ultrasound transducer. PAM: photoacoustic microscopy.

### 2.1 Cascaded gas-filled ARHCF laser

The MIR gas-filled fiber Raman laser source at 3.4 μm is based on a configuration of cascading two gas-filled ARHCFs. To efficiently convert the conventional NIR to the MIR at the selected 3.4 μm wavelength, here the 2^nd^ stage ARHCF is filled with pure H_2_, offering a long vibrational Raman Stokes (VRS) coefficient of 4155 cm^-1^, and also a relatively high gain coefficient compared with other Raman active gases [34]. This requires a pump wavelength at ∼1.4 μm, which is outside the gain range of the known rare-earth ion-based fibers. Therefore, we first generated the required ∼1.4 μm laser by using the 1^st^ order VRS (2331 cm^-1^) of N_2_ in the 1^st^ ARHCF stage [35], pumped by a Yb-doped fiber laser at 1 μm region. The pump laser has an all-fiber structure consisting of a modulated diode laser seed followed by an Yb-doped fiber amplification module, emitting a pulse train with a repetition rate of 1.2 kHz, ∼3.7 ns pulse duration, energy of ∼98 μJ (measured by an energy meter, PE9-ES-C, Ophir Optronics), and 0.128 nm linewidth at 1060 nm wavelength [36]. A λ/2 waveplate is placed at the output of the amplifier for adjusting the polarization orientation to acquire the highest Raman conversion efficiency in the subsequent gas-filled ARHCF system. Then the beam is coupled into the N_2_-filled ARHCF to generate the 1^st^ order VRS line at 1409 nm [35], which is then used as a pump for the 2^nd^ stage ARHCF, to generate the Raman laser at MIR 3.4 μm wavelength.

The scanning electron microscopy (SEM) images of the two ARHCFs used in our experiments are shown in Fig. 1. The first stage ARHCF (Fiber 1 in Fig. 1) is a 16-meters-long single-ring nodeless ARHCF, consisting of 7 rings with a diameter of 16.1 μm and wall thickness of ∼323 nm, forming a negative-curvature core shape with an inner jacket tube diameter of 32.8 μm. The simulated loss spectrum of the fiber is shown in Fig. S1 (a), with a loss value of ∼0.02 dB/m at 1060 nm and ∼0.05 dB/m at 1409 nm [35]. The 2^nd^ stage ARHCF (Fiber 2 in Fig. 1) is 5 meters long with a nested cladding structure and a core diameter of 82 μm [36]. The diameter for external and internal capillaries are 40.3 µm and 13.6 µm, and the wall thickness are 987 nm and 1.37 µm, respectively. This ARHCF has a loss of only ∼0.004 dB/m at 3409 nm (see Fig. S1 (b)), which is a critical condition for the efficient generation of the 3.4 µm Raman laser. In addition, the nested structure of the 2^nd^ ARHCF significantly suppresses the bend loss and therefore allows a bending diameter of ∼40 cm facilitating the development of a compact PAM system. The 1^st^ ARHCF can be also coiled with ∼40 cm diameter without high bending loss due to the small core diameter [37].

### 2.2 Mid-infrared photoacoustic microscopy

The generated 3.4 μm Raman laser beam is then expanded and coupled into the PAM. The output energy is properly attenuated by using a neutral density filter (NDF, Thorlabs, NDC-50C-2M) to avoid the sample damage. The beam is focused by a reflected IR objective lens (PIKE, 40x, 0.78 NA) which has a designed obscuration of 42.9%. The PAM is performed in a transmission mode, and the sample is placed above a transparent sapphire-based coverslip (#18-471, Edmund Optics). Then a water tank with a flat polymer bottom surface (μ-Dish, Ibidi) with 200 μm thickness is placed tightly above the sample to fix and flatten the sample surface and to couple the acoustic signal into the distilled water filled in the tank. The water tank and the sample are combined as an integrated part held by a high-resolution XY stage (8MTF-75LS05, Standa) driven by a stepper & DC motor controller (8SMC5-USB, Standa) to enable the sample scanning with a minimum step size < 10 nm, and a maximum speed of 35000 steps/s. A focused immersive ultrasound transducer (UT, Precision Acoustic) with a central frequency of 20 MHz is immersed within the distilled water in the tank. The UT has an acoustic focal length of 8 mm and a diameter of 10 mm and is coaxially aligned with the reflective IR objective lens. The detected PA signals are filtered by two analog filters (Mini-circuits, 1 MHz long pass filter and 27 MHz low pass filter), then amplified by a low-noise wideband amplifier (Spectrum Instrumentation) and finally received with a high-speed digitizer (M4i.4421-x8, Spectrum Instrumentation) for data processing. The digitizer, integrated in a computer, operates at 250 MS/second sampling rate with a voltage resolution of 16 bits.

An external trigger delay controller (AeroDiode) synchronizes the scanning and data acquisition. A small fraction of the pump pulse energy is extracted from the pump beam and is recorded by a NIR photodetector (DET08C/M, Thorlabs) as a trigger signal, as shown in Fig. 1. Then the output signal from the photodetector is connected to the input side of a trigger delay controller as the input trigger signal. Once the trigger delay controller detects the pump pulse signal, a square signal is generated and acts as the PA trigger for X-Y stage movement and signal synchronization. The PA signal is recorded ∼5 μs after the laser pulse and ∼0.2 μs after the square trigger signal at a repetition rate of 1.2 kHz. During the sample scanning, to minimize noises and enhance signal to noise ratio, 100 pulses (A-lines) are averaged corresponding to a ∼83 ms dwell time.

### 2.3 Brain sample preparation

In order to investigate the performance of our MIR-PAM in real brain tissue, wild-type adult mice were employed. The procedure to prepare the brain slices presented below is approved by the Animal Experiments Inspectorate under the Danish Ministry of Food, Agriculture, and Fisheries, and all procedures adhere to the European guidelines for the care and use of laboratory animals, EU directive 2010/63/EU. A Long-Evans wild-type adult rat was anesthetized by intraperitoneal injection of 200 mg/kg sodium pentobarbital. Hereafter, the level of anesthesia was assessed by pedal reflex pain response to firm toe pinch. Once a sufficient level of anesthesia was reached, the chest cavity was opened to expose the heart. A catheter, connected to a peristaltic pump (Gilsion, minipuls3), was inserted into the left ventricle of the heart, and a small incision was made into the vena cava inferior for perfusion. The perfusion of the rat was performed with 30 ml of 1X phosphate-buffered saline (PBS) followed by 30 ml of 4% paraformaldehyde (PFA) at the rate of 7.8 ml min^−1^. The rat was subsequently decapitated using a guillotine. The brain was dissected out and directly put into ice-cold 4% PFA. After fixation for 4 h at 4 °C, the brain was transferred to 30% sucrose (w/v) for cryoprotection at 4 °C. A vibratome (LEICA, VT1200) was then used to section the brain into slices. To facilitate sectioning, the brain was embedded in 3% agarose gels prepared with a method similar to the one for preparing the 0.6% gels described above. The slicing was conducted in ice-cold 1X PBS. A similar process has been also followed in [38]. The brain slice has a thickness of 400 μm. During the PAM imaging, the slice is kept in 1X PBS at room temperature.

## 3 Results and discussion

### 3.1 Gas-filled hollow-core fiber laser source

Figure 2 (a) presents the spectra of the pump laser as well as Raman Stokes lines, measured using an infrared spectrometer (Spectro320 Instrument Systems) with a resolution of 0.14 nm. The gaussian-like beam profile of the 1^st^ Raman pulse, measured by a beam profiler (BP109-IR2, Thorlabs), indicates that the Raman laser operates at the fundamental mode. To maximize the output pulse energy, first, we measured the pulse energy of the 1409 nm Raman line output of the 1st stage ARHCF in terms of the N_2_ pressure [35]. The Raman line appears at 9 bar pressure and the pulse energy reaches its maximum of up to ∼26.5 µJ at ∼15 bar pressure, corresponding to a quantum efficiency of 45%.

**Fig. 2.**
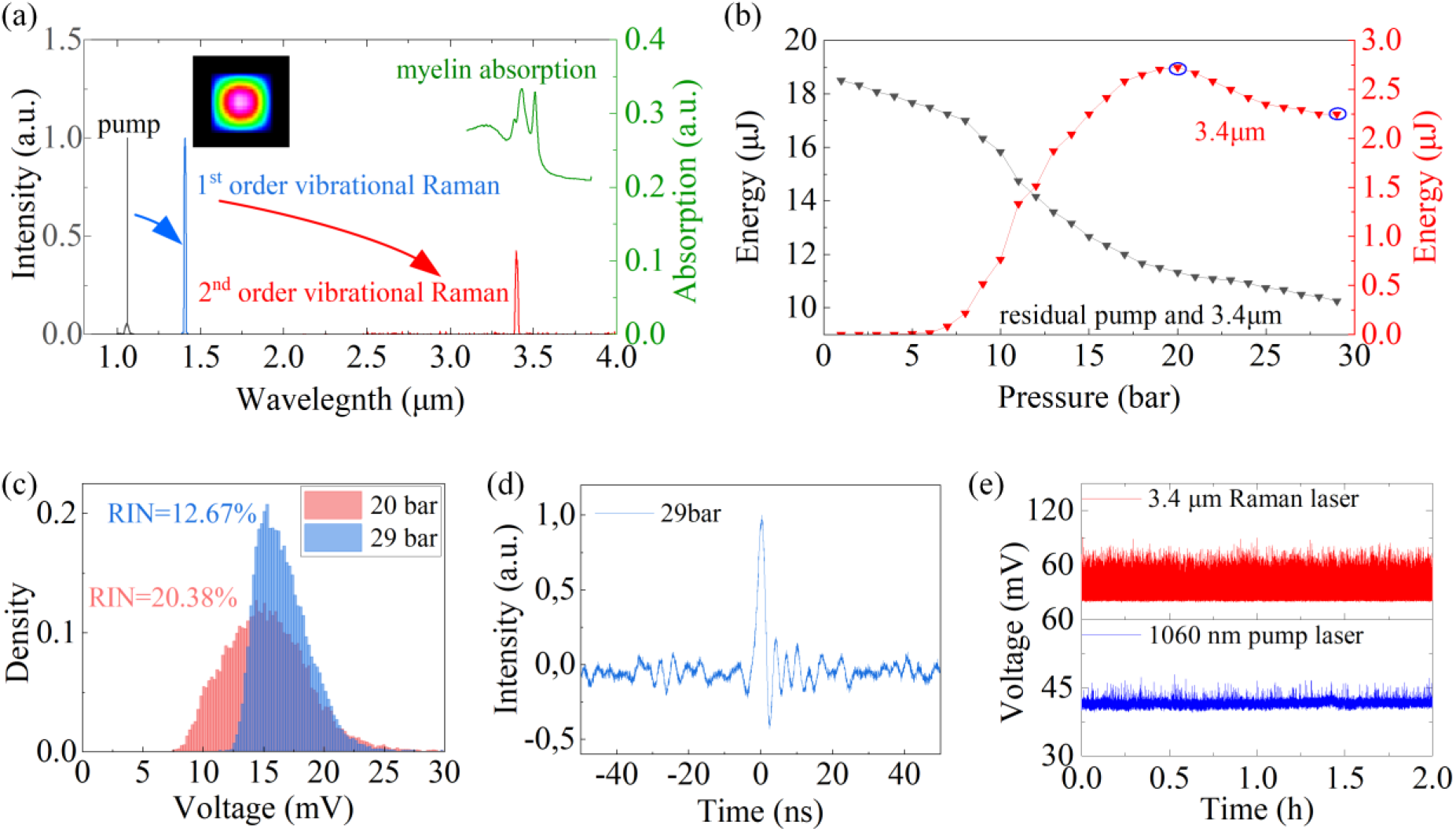
Characterization of the gas-filled ARHCF laser source. **(a)** Measured spectra including the pump line and Raman lines generated from the cascaded ARHCFs. The right axis shows the absorption spectrum of myelin extracted from [10]. Inset: beam profile of 1^st^ Raman pulse at 1409nm. **(b)** Pulse energy evolution of the 3.4 μm Raman pulse as a function of H_2_ pressure. **(c)** Histograms of the pulse peak intensity of the 3.4 μm Raman pulse at 20 bar and 29 bar, respectively. **(d)** Pulse profile of the 3.4 μm Raman laser. **(e)** Pulse peak intensity monitoring of the 1060 nm pump and the 3.4 μm Raman lasers over 2 hours.

By pumping the 1409 nm Raman line into the 2^nd^ stage ARHCF, a Raman line at 3.4 μm wavelength is observed when the pressure is > 5 bar, as seen in Fig. 2 (a), with a linewidth of ∼1 nm. The average power of the 3.4 μm Raman laser was measured by extracting it from the residual 1.4 μm pump using a 2400 nm long-pass filter (FELH2400, Thorlabs, transmission 98% at 3.4 μm). Figure 2 (b) shows the pulse energy evolution of the 3.4 μm Raman laser, as well as the total energy including both the Raman laser and residual pump at 1409 nm without using a filter, as a function of the H_2_ pressure. The Raman pulse energy begins to rise at ∼5 bar, with the highest value to be 2.75 μJ at ∼20 bar. As the pressure further increases, the energy starts to decrease slightly, because the Raman laser reaches its maximal pulse energy at a shorter fiber length. The long-term stability and noise performance of the Raman laser source were also characterized, to provide a reference for implementing the subsequent PAM application. The noise performance of the MIR Raman pulses was measured in term of the relative intensity noise (RIN) of pulse peak intensity at 20 and 29 bar, using a photodetector (100 MHz bandwidth, PDAVJ8, Thorlabs) connected to an oscilloscope (6 GHz bandwidth, MSO64B, Tektronix). Figure 2 (c) shows the distribution of the measured histograms. At 20 bar pressure, the distribution has a gaussian-like profile with a RIN of 20.38%. Compared to 20 bar, at 29 bar pressure although the pulse energy slightly decreases to 2.25 μJ, a lower RIN of 12.67% is obtained. This is because, the SRS process becomes more efficient towards higher H_2_ pressure due to the further suppression of transient SRS regime [39]. Given this property, here we set the pressure of H_2_ to 29 bar, to better mitigate the acoustic signal fluctuation in the PAM application. Figure 2 (d) shows a typical pulse profile of the 3.4 µm Raman laser. The pulse width is ∼2 ns but the precision of this measurement is compromised because of the limited bandwidth of the MIR photodetector. This Raman laser also shows a good long-term stability, as indicated by the 2 hours monitoring results of the laser peak intensities of both the 3.4 μm Raman laser and its pump at 1060 nm.

### 3.2 Characterization of the MIR-PAM

The lateral resolution of the PAM was first evaluated by imaging a sharp knife edge with a thickness of 100 μm. Figure 3 (a) shows the optical image (top) and the corresponding PAM image (bottom). The pulse energy before and after the objective lens are ∼1.9 μJ and ∼1 μJ respectively due to the 42.9% obscuration. The PAM images are obtained with a step size of 1.25 μm after averaging 100 pulses. The edge spread function (ESF) is acquired from the averaged raw data of the intensity, and the line spread function (LSF) is then calculated and fitted from the differential of the ESF, as shown in Fig. 3 (b). From the full width at half maximum (FWHM) of LSF, the lateral resolution of our MIR-PAM is estimated to be ∼12.36 μm.

**Fig. 3.**
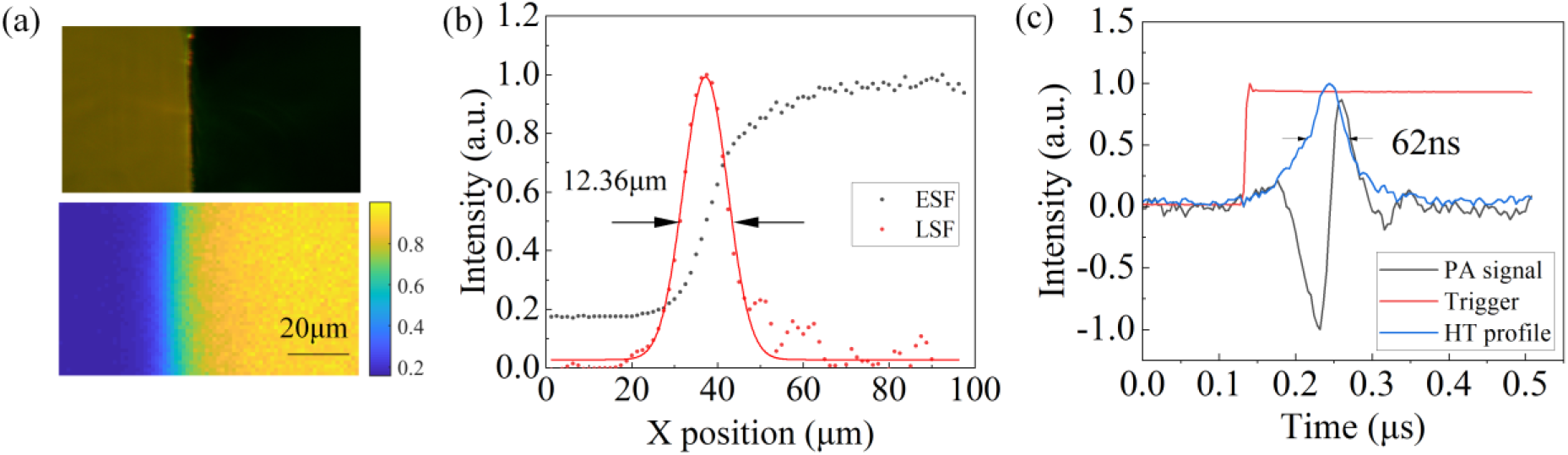
Resolution characterization of the MIR-PAM. **(a)** Optical and photoacoustic images of a sharp blade edge. **(b)** Fitted ESF and LSF extracted from PA image in (a). **(c)** Single PA signal and its Hilbert transformation (HT) profile.

The axial resolution *R*_*a*_ is estimated experimentally by the pulse width of the impulse response of a single PA signal. Figure 3 (c) shows the PA signal and its envelope after Hilbert Transformation (HT). The axial resolution is extracted from the FWHM of the envelope, which is 62 ns, corresponding to 95.5 µm in distance, given the propagation velocity of sound in the sample of *c*=1540 m/s [40]. The axial resolution can also be numerically calculated by the velocity of the acoustic signal and the central frequency of the ultrasound transducer as *R*_*a*_=0.88*c*/*B*, where *B* is the central frequency of the UT. In our case, the theoretical axial resolution is ∼67.8 μm.

### 3.3 Ex-vivo PAM

Figure 4 (a) shows the *ex-vivo* PAM image of the mouse brain slice with the thickness of 400 μm obtained through using the proposed 3.4 μm laser as light source. The scanning step size is 40 μm and the total scanning time is ∼40 min. HT was used for extracting the absolute amplitude of the acoustic signal, and then the image is post-processed by Gamma transformation and a contrast enhancement of 0.5% saturation using ImageJ software. In comparison with the optical image in Fig. 4 (b), we can clearly see the outlines of the different brain regions. The myelin-rich regions, such as corpus callosum, are brighter than other regions. Small structures like mammillothalamic fasciculus (mtf) and fornix (fo) can also be distinguished, exhibiting as bright spots, since they consist of heavily myelinated fibers [10,41,42]. These characteristics are invisible in the optical image shown in Fig. 4 (b). The regions with less myelin, such as hippocampus, which mainly consists of gray matter, appear to be darker. Finally, due to the lack of myelin, the cavity structure (lateral ventricle IIId) looks like a void with clear boundaries.

**Fig. 4.**
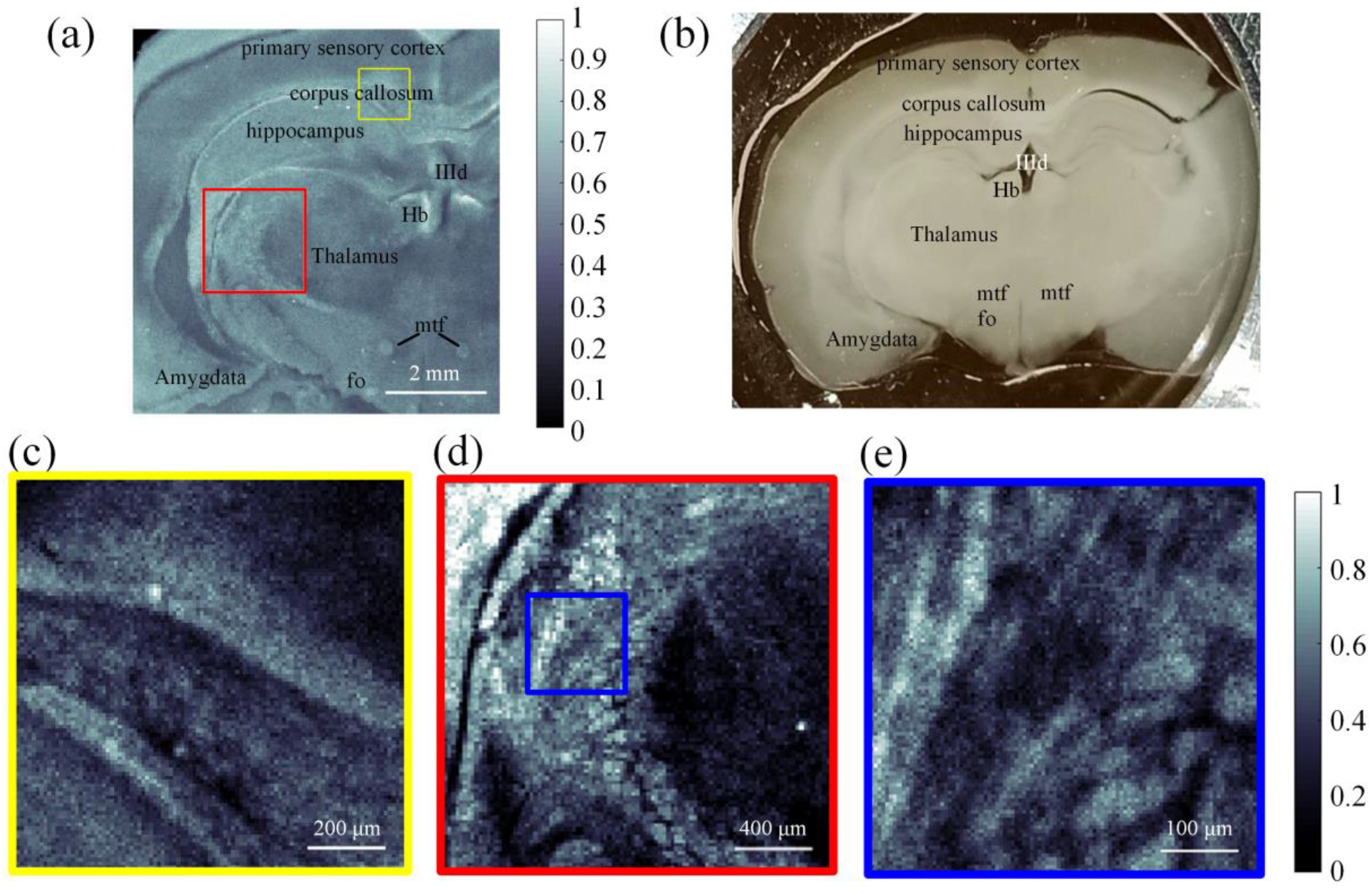
Image of the mouse brain. **(a)** *Ex-vivo* PA image of the mouse brain slice. Hb: habenular nuclei. IIId: IIIrd ventricle. mtf: mammillothalamic fasciculus. fo: fornix. **(b)** Optical image of the mouse brain slice. **(c)** Enlarged view of the structure in the yellow box in (a) with a scanning step size of 10 μm. **(d)** Enlarged view of the structure in the red box in (a) with a scanning step size of 20 μm. **(e)** Enlarged view of the structure in the blue box in (d) with a scanning step size of 5 μm.

Enlarged views of the corpus callosum and thalamus regions were scanned with a step size of 10 μm and 20 µm respectively, as shown in Fig. 4 (c) and (d). The border between corpus callosum and hippocampus is clearly visible in Fig. 4 (c). In Fig. 4 (d), patterns with a fish-scale-like appearance are observable, which agree with other brain images reported in the literature [10]. A further enlargement of the thalamus region, performed with a scanning step of 5 µm, is shown in Fig. 4 (e). Here, we can observe some thick nerve fiber bundles, which has been also verified and observed by other MIR-PAMs [10,13], demonstrating the ability of our new MIR-PAM Raman fiber laser for lipid-rich myelin imaging. All enlarged images required a scanning time of about 20 min.

## 4 Conclusion

This work aims to create a new avenue of the emerging gas-filled Raman laser technology for MIR-PAM. As a proof-of-concept, a high-energy Raman laser at ∼3.4 µm was developed and employed for brain imaging. This work can be further extended to multispectral PAM through operating the gas-filled fiber laser with the reconfigurable multiple spectral lines spanning from near-to mid-infrared region [36].

On the other hand, although this work still involves the free-spacing coupling to ARHCF, an all-fiber structure will be expected in our future work, given the recent progress on the low-loss splicing of ARHCFs [43-45].

### Disclosures

The authors declare that there are no conflicts of interest.

### Code, Data, and Materials

The data of this work is available from the corresponding author upon reasonable request.

## Acknowledgments

This work is supported by LUNDBECK FONDEN (Grant No. R346-2020-1924, No. R276-2018-869), VILLUM Fonden (Grant No. 36063, Grant No. 40964), and US ARO (Grant No. W911NF-19-1-0426).

## Supplementary Material

### SUPPLEMENTARY NOTE 1. State-of-the-art of MIR gas-filled anti-resonant hollow-core fiber (ARHCF) lasers

**Table 1.**
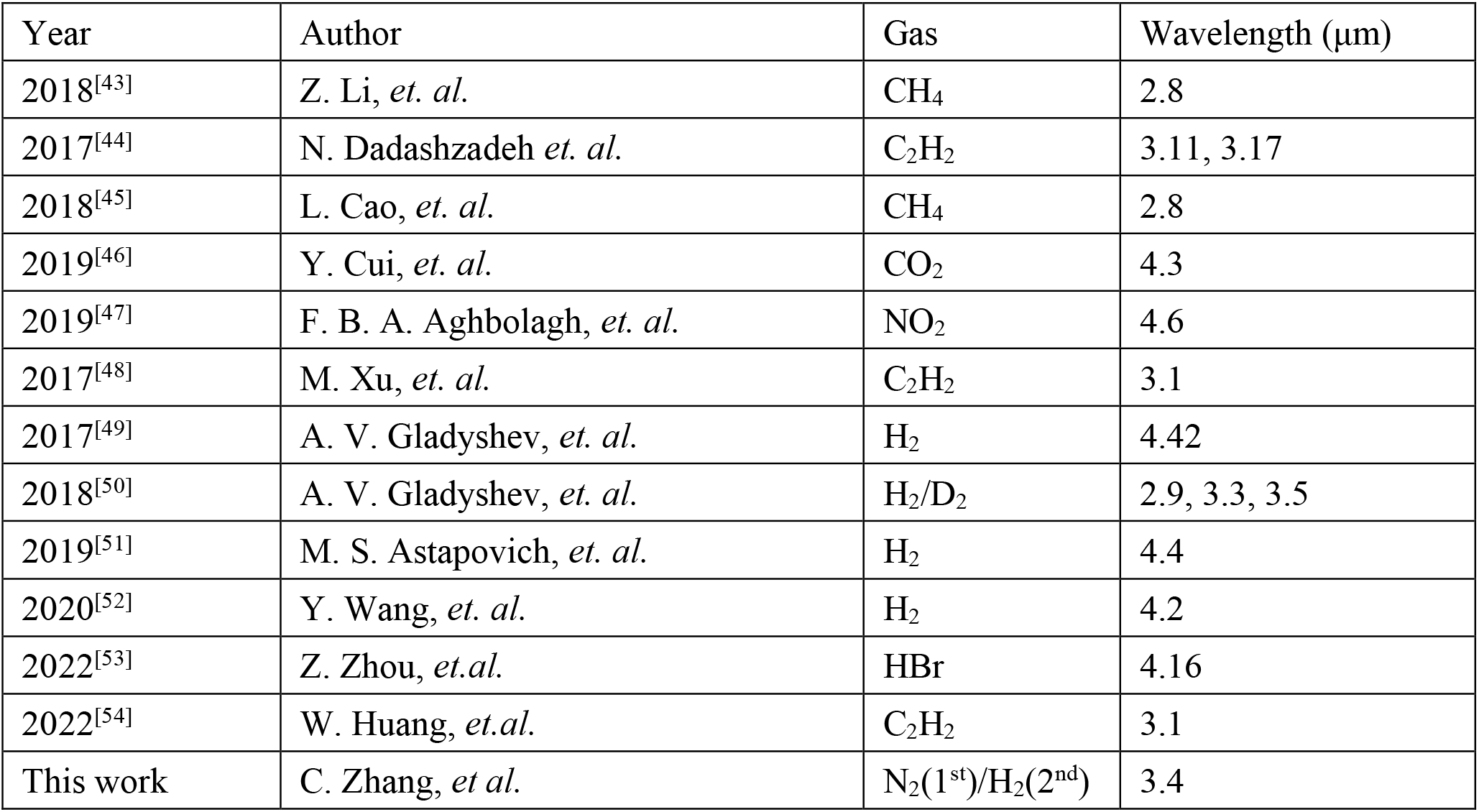
State-of-the-art of MIR gas-filled ARHCF lasers.

| Year | Author | Gas | Wavelength ($\mu\text{m}$) |
| --- | --- | --- | --- |
| 2018<sup>[43]</sup> | Z. Li, *et. al.* | CH<sub>4</sub> | 2.8 |
| 2017<sup>[44]</sup> | N. Dadashzadeh *et. al.* | C<sub>2</sub>H<sub>2</sub> | 3.11, 3.17 |
| 2018<sup>[45]</sup> | L. Cao, *et. al.* | CH<sub>4</sub> | 2.8 |
| 2019<sup>[46]</sup> | Y. Cui, *et. al.* | CO<sub>2</sub> | 4.3 |
| 2019<sup>[47]</sup> | F. B. A. Aghbolagh, *et. al.* | NO<sub>2</sub> | 4.6 |
| 2017<sup>[48]</sup> | M. Xu, *et. al.* | C<sub>2</sub>H<sub>2</sub> | 3.1 |
| 2017<sup>[49]</sup> | A. V. Gladyshev, *et. al.* | H<sub>2</sub> | 4.42 |
| 2018<sup>[50]</sup> | A. V. Gladyshev, *et. al.* | H<sub>2</sub>/D<sub>2</sub> | 2.9, 3.3, 3.5 |
| 2019<sup>[51]</sup> | M. S. Astapovich, *et. al.* | H<sub>2</sub> | 4.4 |
| 2020<sup>[52]</sup> | Y. Wang, *et. al.* | H<sub>2</sub> | 4.2 |
| 2022<sup>[53]</sup> | Z. Zhou, *et.al.* | HBr | 4.16 |
| 2022<sup>[54]</sup> | W. Huang, *et.al.* | C<sub>2</sub>H<sub>2</sub> | 3.1 |
| This work | C. Zhang, *et. al.* | N<sub>2</sub>(1<sup>st</sup>)/H<sub>2</sub>(2<sup>nd</sup>) | 3.4 |

### SUPPLEMENTARY NOTE 2. Loss spectrum of ARHCFs

**Fig. S1.**
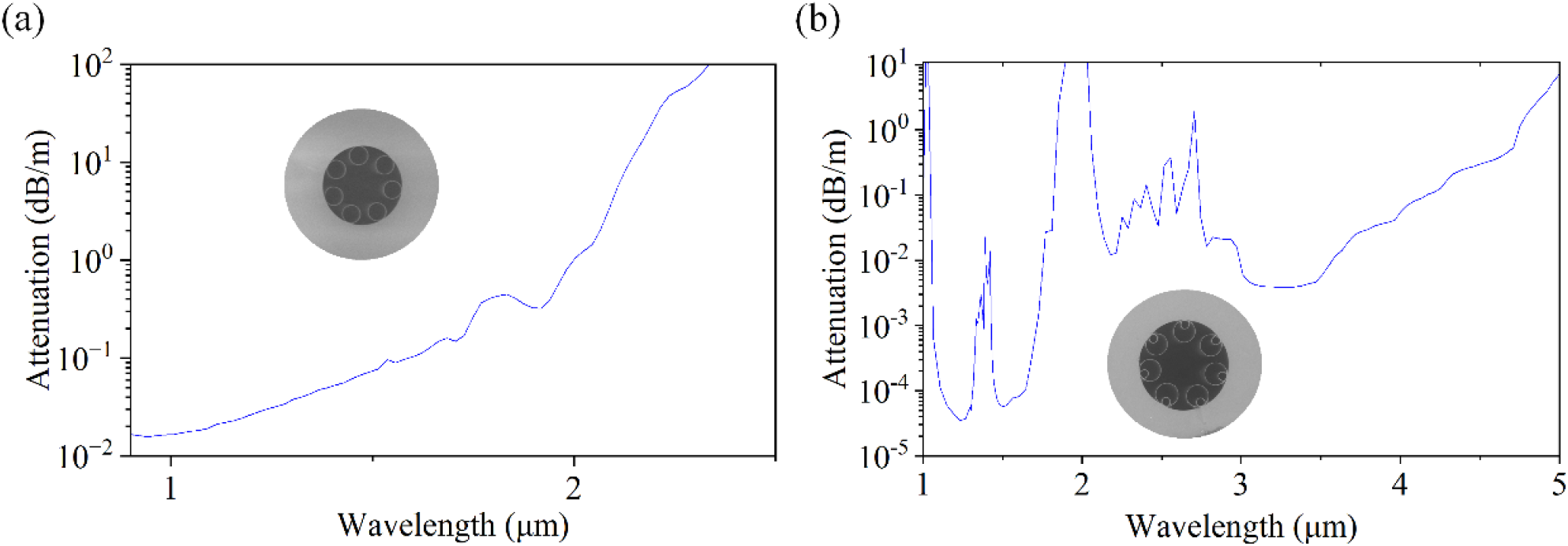
Simulated attenuation spectra of **(a)** the first stage ARHCF and **(b)** the second stage ARHCF.

For the first stage ARHCF, the simulated attenuation coefficient is ∼0.02 dB/m at 1060 nm and ∼0.05 dB/m at 1409 nm, respectively. For the second stage ARHCF, the attenuation coefficient is ∼0.005 dB/m at 1409 nm and ∼0.004 dB/m at 3.4 μm, respectively.

